# Challenges in antibody structure prediction

**DOI:** 10.1101/2022.11.09.515600

**Authors:** Monica L. Fernández-Quintero, Janik Kokot, Franz Waibl, Anna-Lena M. Fischer, Patrick K. Quoika, Charlotte M. Deane, Klaus R. Liedl

**Author notes:** These authors contributed equally (Monica L. Fernández-Quintero, Janik Kokot, Franz Waibl).

## Abstract

The tremendous advances in structural biology and the exponential increase of high-quality experimental structures available in the PDB motivated numerous studies to tackle the grand challenge of predicting protein structures. AlphaFold2 revolutionized the field of protein structure prediction, by combining artificial intelligence with evolutionary information.

Antibodies are one of the most important classes of biotherapeutic proteins. Accurate structure models are a prerequisite to advance biophysical property predictions and consequently antibody design. Various specialized tools are available to predict antibody structures based on different principles and profiting from current advances in protein structure prediction based on artificial intelligence. Here, we want to emphasize the importance of reliable protein structure models and highlight the enormous advances in the field. At the same time, we want to raise the awareness that protein structure models—and in particular antibody models—may suffer from structural inaccuracies, namely incorrect cis-amid bonds, wrong stereochemistry or clashes. We show that these inaccuracies affect biophysical property predictions such as surface hydrophobicity. Thus, we stress the significance of carefully reviewing protein structure models before investing further computing power and setting up experiments. To facilitate the assessment of model quality, we provide a tool “TopModel” to validate structure models.

## Breakthroughs in Protein/Antibody Structure Prediction

Predicting the three-dimensional (3D) structure of a protein based solely on the amino-acid sequence has been one of the grand challenges in the field of protein structure prediction.^1^ Accurate prediction of the 3D structure of a protein is critical to understand its function, as the shape of the protein determines its properties and ultimately its function. To determine/identify the state-of-the-art methods in protein structure prediction, the biennial community-based benchmarking experiment “Critical Assessment of methods in protein Structure Prediction (CASP)” has been established.^2–4^ In CASP14 (2020), DeepMind showcased AlphaFold2, which is a program based on artificial intelligence (AI) that directly processes multiple sequence alignments.^5^ Comparable accuracies in predicting protein structures could also be achieved with RoseTTAFold^6^, and specialized tools for antibodies which incorporate the recent advances have been presented.^7–9^ Those tools are highly accurate based on global measures, often with RMSDs to the crystal structure of less than 1 Å. However, there might be higher inaccuracies in parts of the protein. Post-translational modifications are omitted, but can sometimes be added afterwards.^10^ Furthermore, the accuracy for multimers, such as antibodies, is still lower.^11^ Additional challenges can arise for antibodies since VDJ recombination events do not follow the classical pathway of evolution.^12^

Antibodies are crucial components of the adaptive immune response.^13^ Genetic recombination and somatic hypermutation events enable the adaptive immune system to produce a vast number of antibodies against a variety of pathogens.^12^ To understand and optimize antigen recognition and to enable rational design of antibodies, accurate structure models are essential.^14^ Despite these recent advances, accurate structure prediction of antibodies remains challenging and still needs to be extensively validated. In particular, the flexible loops involved in recognizing the antigen pose a major challenge.^15,16^ In comparison to other protein superfamilies, the fold of antibodies is generally highly conserved.^17–19^ Especially the framework of the antigen-binding fragment (Fab) is structurally almost identical for all antibodies.^20,21^ However, the area hardest to predict accurately is the six hypervariable loops that form the antigen-binding site, called the paratope. These loops are also known as the complementarity determining region (CDR) and provide the structural diversity essential to recognize a wide range of antigens. Five of the six loops tend to adopt canonical cluster folds based on their length and sequence composition. However, the third CDR loop of the heavy chain, the CDR-H3 loop, is the most diverse in length, sequence and structure and therefore is the most challenging loop to predict accurately. Additionally, the CDR-H3 loop conformation is also strongly influenced by the relative interdomain orientation, as it is located in the center, directly in the interface between the heavy and the light chains. Recently, to tackle this challenge, various antibody-specific deep learning methods such as ABlooper, DeepAb and IgFold have significantly improved the CDR loop modeling accuracy.^7– 9,22^ The predicted structure models achieve similar or better quality than methods, that are able to predict all types of protein structures (including AlphaFold2).^11,23^ All these improvements have enabled predictions of a vast number of antibody structures at a high level of accuracy, which can then further be used as input structures for virtual screening or to inform rational design of antibodies.^14,15^

### Possible inaccuracies in antibody structure models

Here, we investigated a dataset consisting of 137 antibody sequences published by Jain et al. and used several freely available antibody structure prediction tools, i.e., ABlooper^7^, IgFold^8^, DeepAb^9^ and the MOE Antibody Modeller^24^, to generate structure models for further biophysical characterizations.^25,26^ Careful inspection of the generated models revealed inconsistencies such as cis-amid bonds in the CDR loops, D-amino acids and severe clashes. In total, this resulted in up to 300 D-amino acids and up to 240 cis-amid bonds for the 137 antibody models. We found cis-amid bonds and clashes independent of the applied antibody modelling tools. The only tool that did not introduce any D-amino acids is DeepAb. These structural inaccuracies affect the results of structure-based biophysical property predictions.^25,26^ In addition to the Jain et al. dataset^27^, we predicted the structure of the CIS43 antibody, where the experimental X-ray structure (PDB accession code: 7SG5)^28^ was released after the structure prediction tools were published, to have an experimental reference structure outside of the used training set. Figure 1 shows an overlay of all obtained structure models and reveals an overall high structural similarity, reflected in low overall RMSD values (∼1Å). However, substantial structural variability can be observed in the CDR-H3 loop (RMSD values >2Å). This result points out one particular challenge in predicting antibody structures: The high variability of the antibody CDR loops cannot be captured/represented by one single-static structure. While there can be properties and metrics that are not too much affected by these issues, metrics that rely on accurate CDR-H3 structures will be strongly distorted. This includes antibody-antigen docking, since the CDR-H3 loop is a central part of the binding interface, as well as structure-based hydrophobicity calculations, since the binding interface frequently contains more hydrophobic amino acids than the rest of the antibody surface.^14,15,25^ Molecular dynamics simulations (MD) might correctly capture the ensemble in solution if the starting structure is sufficiently close, but current MD approaches cannot sample cis-trans isomerisation or transitions from D-to L-amino acids, leading to ensembles that are potentially worse than the starting structure in terms of RMSD to the crystal structure (SI Figure S1).

**Figure 1:**
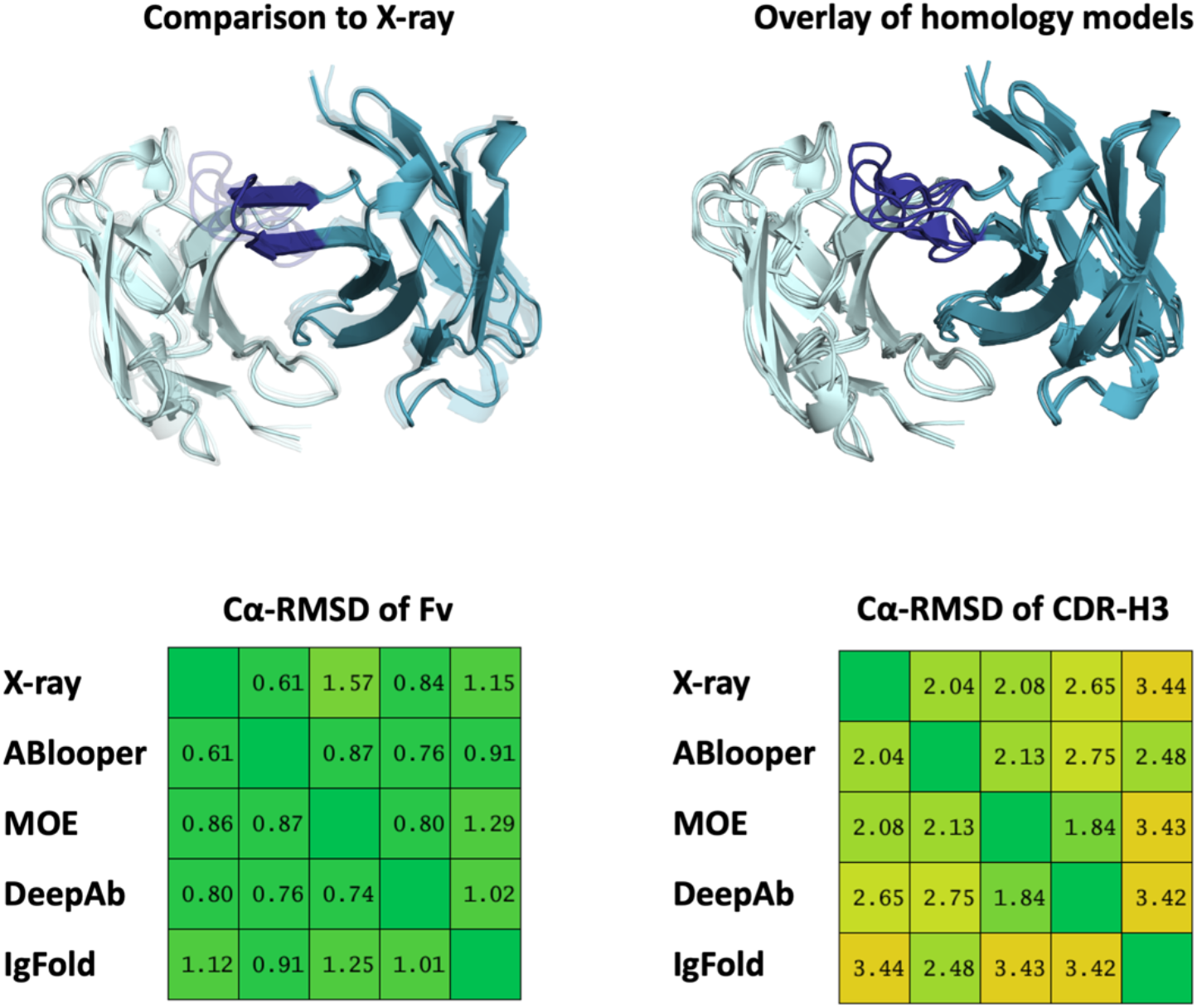
A) Comparison of the available X-ray structure of the CIS43 antibody (PDB code: 7SG5) with the structure models generated with different antibody prediction tools, namely ABlooper, MOE, DeepAb and IgFold. B) Structural overlay of the obtained Fv models, showing the high variability in the CDR-H3 loop. C) Cα-RMSD matrix of the X-ray structure and the respective models for the whole Fv. D) Cα-RMSD matrix of the X-ray structure and the respective models for the CDR-H3 loop.

To show the effect of starting structures with cis-amid bonds and D-amino acids in the CDR loops, we compare the surface hydrophobicity of structure models for the CIS43 antibody variant with the X-ray structure (Figure 2). The surface hydrophobicity was assigned using the hydrophobicity scale by Wimley and White.^29^ We find differences in the surface hydrophobicity, which is expected as hydrophobicity is potentially a strongly conformation dependent property, since small sidechain rearrangements may expose otherwise buried hydrophobic groups. While small inaccuracies in the atomic positions can be fixed by molecular dynamics simulations, the correct sidechain packing is often impossible when D-amino acids or cis-amide bonds are present, leading to almost irreparable errors in the biophysical property estimation. The same is true for antibody-antigen docking, where an accurate representation of the surface is required to find the correct interactions with the antigen.

**Figure 2:**
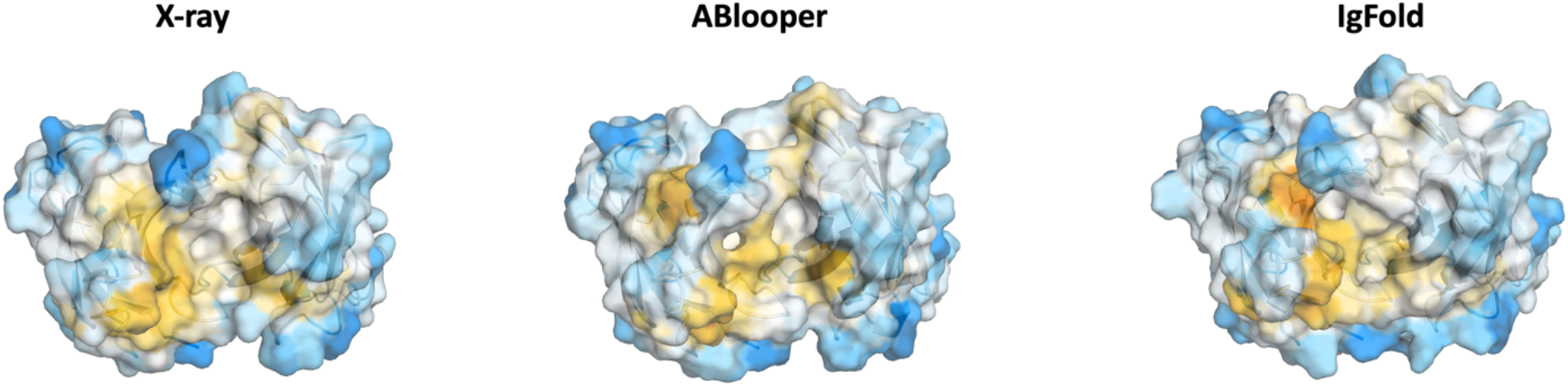
Surface hydrophobicity mapped on the X-ray structure and two antibody models. Surface hydrophobicity was assigned by the Whimley and White hydrophobicity scale. Hydrophobic areas are colored in yellow, while hydrophilic parts are depicted in blue.

Figure 3 shows examples of cis-amide bonds and D-amino acids in the CDR-H3 loop of CIS43. Additionally, in one of the obtained models we find a missing proline sidechain at the tip of the CDR-H3 loop. Rebuilding the proline results in severe clashes, clashes, as shown in the right panel of Figure 3, and the only way to avoid these clashes would be to build a D-Proline instead.

**Figure 3:**
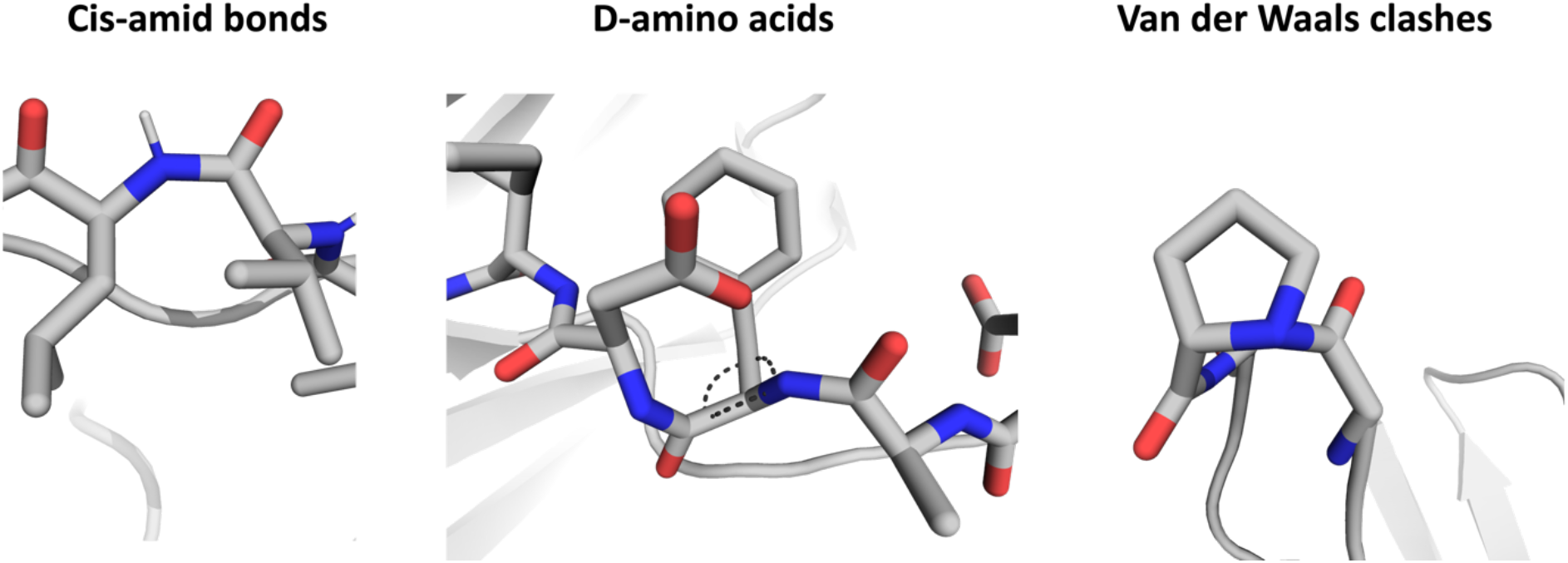
Examples of structural inaccuracies observed in some of the models, namely cis-amid bonds, D-amino acids and Van der Waals clashes.

To facilitate the identification of these issues, we present a tool (available on Github: https://github.com/liedllab/TopModel), called “TopModel”, that quickly checks the structure for cis-amid bonds, D-amino acids, and clashes. With this tool, structure models can rapidly be checked to assess the quality/accuracy of the models before performing further analysis. At the same time, it offers the possibility to directly visualize these issues in PyMOL.^30^ Figure 4 shows the output obtained by “TopModel”. Residues colored in magenta (D-amino acids) and red (cis-amid bonds) represent issues that should be fixed, while cis-Prolines are colored green, as they occasionally occur in native protein structures. Van der Waals clashes are colored in yellow. For the clashes we also provide an optional score, to quantify the quality of the model, that takes the number of clashes and the length of the protein into account, to quantify the quality of the model. Non-planar amid bonds are depicted in cyan. As these issues can also be found for other protein structure models apart from antibodies, we recommend checking every model with “TopModel” and stress the importance of validating the obtained structures to ensure the most accurate results/predictions as possible.

**Figure 4:**
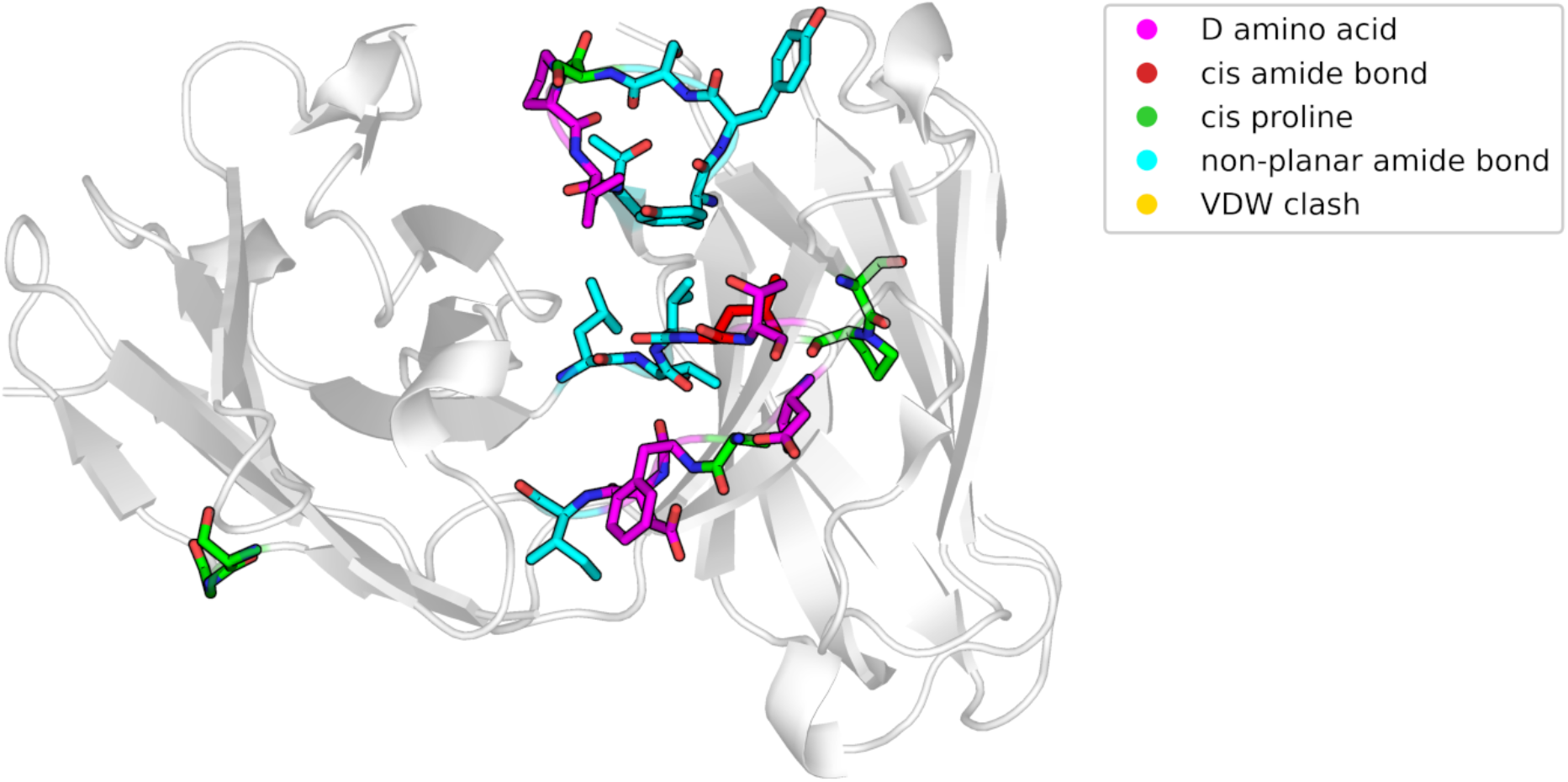
Pymol visualization of the structural inaccuracies, as output of the “TopModel” tool.

## Discussion

While we want to highlight the enormous advances in the protein structure prediction field, at the same time, we want to emphasize the importance to critically review structure models--especially antibody models--before basing conclusions/further experiments/molecular dynamics simulations on potentially erroneous models.^31^ In particular, we want to stress the importance of not limiting the characterization of the antigen binding site to a single-static structure model, as there is already a high structural variability between the different models. This high variability suggests that even considering ensembles in solution might more accurately reflect the properties/functions of the antibody.^16^ The tremendous development in protein structure prediction enables fast protein structure predictions that approach the accuracy of experimental structures.^23^ These breakthroughs have been achieved by combining artificial intelligence with an effective exploitation of the available structural information and incorporation of evolutionary related sequences in terms of multiple sequence alignments (MSAs).^23^ Thereby, AlphaFold, RoseTTAFold and ESMFold revolutionized protein structure prediction.^6,23,32–34^ Various deep learning (based) approaches have also been shown to improve antibody structure prediction and outperform all previously available antibody structure prediction methods.^7–9,35^ Accurately predicting the structure of antibodies is central to understand their function, to elucidate antibody-antigen binding and inform rational antibody design.

However, the most challenging part in antibody structures prediction is concentrated in the six CDR loops, as they reveal the highest variability in both sequence and structure.^13,36^ In particular, the CDR-H3 loop reveals the highest diversity and variability, which impedes the accurate prediction of its structure.^37,38^ This is in line with our findings, as the comparison of different antibody prediction methods reveals the most diverging results for the CDR-H3 loop. Figure 1 shows an overlay of all the obtained models, highlighting the high conformational variability of the CDR-H3 loop. To account for this high diversity of the CDR loops one single static structure might not be sufficient and therefore the CDR loops should rather be characterized as ensembles in solution.^16,37^ This is especially important, as various biophysical properties of antibodies, such as hydrophobicity, are conformation dependent and already small sidechain rearrangements reveal distinct surface properties.^25,26^ Molecular dynamics simulations provide such ensembles in solution, increasing the probability that conformations determining biophysical properties are captured.^25^ In agreement with these observations, we find that these conformational differences between the models can result in changes in the surface hydrophobicity.

In addition to the high divergence in CDR-H3 loop conformations, we found various cis-amid bonds, D-amino acids, and clashes in the obtained models. Such modeling artifacts are firstly non-natural and secondly, can strongly influence biophysical property predictions and result in misleading conclusions.^31^ Thus, to address these pitfalls, we provide a tool that quickly inspects protein structure models and identifies issues/flaws in the protein structures, namely the python package “TopModel”. As accurate antibody structures are a prerequisite to reliably understand antibody function and characterize biophysical properties, we strongly suggest an additional validation of the respective structure models to increase the quality of the respective predictions.

## Methods

The tool “TopModel” (version 1.0) inspects and highlights issues in a structure model. “TopModel” checks the chirality, amide bond stereochemistry and Van der Waals clashes for every residue in the structure model. The structure models are parsed and analyzed using biopython.^39^ To calculate the chirality a triangle is defined based on the priority of the atom chains around the chiral center. The direction of a normal vector to this plane, calculated using the cross product, is determined by the priority of the atom chains. By calculating the dot product of this normal vector and a vector from the chiral center to the plane the orientation of the three atom chains with respect to the center can be determined. Based on the sign of the resulting scalar the chirality can be assigned. This approach allows us to calculate the chirality even if no hydrogens are included in the structure model. The amide bond is inspected by calculating the dihedral angle of the protein backbone. Cis-amide bonds to proline are labelled separately as they naturally occur more frequently than in other amino acids. Dihedral angles that could neither be assigned cis nor trans are labelled as non-planar. The VDW clashes are quickly computed using a k-d tree^40^ and VDW radii data gathered using the python package mendeleev^41^. All pairs of atoms within 5 Å of each other are checked for VDW clashes by calculating the distance and comparing against the combined VDW radii minus 0.5 Å. The pairs of atoms that are closer than 5 Å are calculated using a k-d tree^40^ and the VDW radii data is gathered using the python package mendeleev^41^.For the clashes an optional score can be displayed, which takes the number of clashes and the length of the protein into account. The chiralities and amide bond orientations are not included in the score. To quickly assess the structural implications of the issues found by “TopModel”, the analyzed structure can be opened in PyMOL^30^ with the issues highlighted and labeled as shown in Figure 4.

## Author Contributions

M.L.F.Q., J.K., F.W. performed research, analyzed data and drafted the manuscript. A.M.F. performed research and analyzed data. P.K.Q. analyzed data and contributed in writing the manuscript. C.M.D. and K.R.L. supervised the research.

## Acknowledgement

The computational results presented her have been achieved (in part) using the Vienna Scientific Cluster (VSC). This work was supported by the Austrian Science Fund (FWF) under grant number P34518. MFQ received the APART-MINT PostDoc fellowship of the Austrian Academy of sciences (No. 11985). We acknowledge PRACE for awarding us access to Piz Daint at CSCS, Switzerland.

