## Supporting Information for "Challenges in antibody structure prediction"

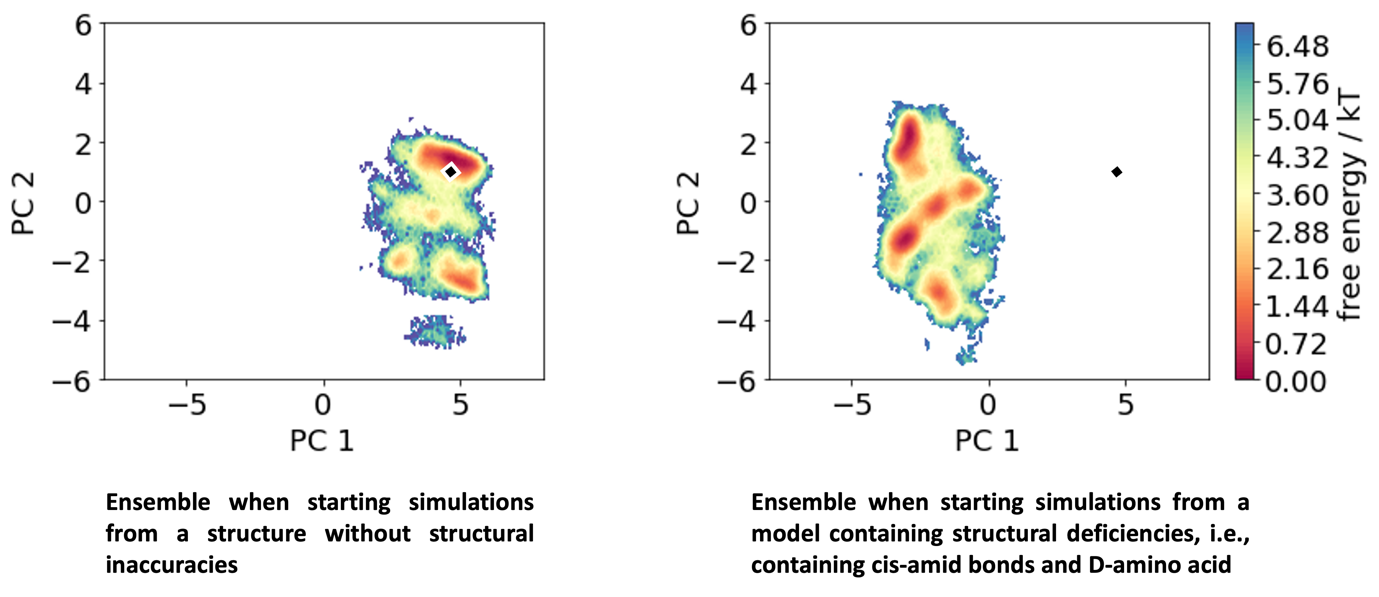

SI Figure S1: Principal Component Analysis (PCA) comparing an MD ensemble in solution starting from a structure without structural inaccuracies (left) with an ensemble obtained using a structure with deficiencies. The diamond projected into the PCA space represents the available X-ray structure for the CIS43 antibody (PDB accession code: 7SG5).
